# Establishment of an avian influenza surveillance program in Australia’s largest river basin

**DOI:** 10.64898/2026.09.09.750291

**Authors:** MV Jackson, HM McGinness, M Wille, M Davies, A Read, B Farrugia, R Doble

## Abstract

To-date Australia has had mostly coastal occurrences in a small but growing number of species of high pathogenicity avian influenza (HPAI) H5N1 clade 2.3.4.4b. The potential impacts of expected spread are the focus of significant preparation activities. Here we report on early surveillance undertaken in 2025-26 in the Murray-Darling Basin, Australia’s largest river system, which contains internationally important wetlands that support large multi-species aggregations of waterbirds vulnerable to mass mortality. We detected two low pathogenicity avian influenza (LPAI) viruses of Australian origin in wild waterbirds. Ongoing surveillance will aid in early detection and rapid response in the case of a major inland outbreak of H5N1.

## Main text

Since emerging in Europe in 2021, high pathogenicity avian influenza (HPAI) H5N1 clade 2.3.4.4b (hereafter H5 bird flu) has spread throughout the world causing a panzootic. It has been detected in >400 bird species (Ryding et al. 2026) and caused mass mortality in a wide range of wild bird and marine mammal species with severe population declines in seabird species in multiple regions (Knief et al. 2026). It has also caused damage to the poultry industry (Krammer et al. 2025), dairy industry (Rodriguez et al. 2025), and caused human cases (Garg et al. 2025).

Given the global spread of H5 bird flu, it was only a matter of time until viral incursion to the Australian continent occurred. Each year millions of shorebirds migrate into Australia from breeding grounds in the northern hemisphere through regions with active outbreaks. Given this high-risk pathway, targeted virus surveillance of migratory seabirds and shorebirds has been undertaken in coastal areas of Australia since 2022 (Wille et al. 2026a; Wille et al. 2026b). A diversity of low pathogenicity avian influenza viruses was detected in shorebirds in 2024 and 2025, underscoring the potential for incursion through long-distance migrants (Wille et al. 2026a; Wille et al. 2026b). Further, Australia’s sub-Antarctic islands host considerable populations of pelagic seabirds and marine mammals, all of which travel considerable distances. Indeed, H5 bird flu was confirmed on the Australian sub-Antarctic Island of Heard Island in Oct 2025 (McInnes et al. 2026), and incursion to Australia via this southern route occurred in June 2026. To-date there have been mainland occurrences of H5N1 mostly in coastal areas (but with some limited inland incursions) in a small but growing number of species, mostly seabirds but also some mammals.

In addition to migratory shorebirds and seabirds, Australia is home to many other highly mobile waterbird species that could both facilitate virus spread and experience mass mortalities. For example, nomadic and partially migratory ducks and group-nesting waterbirds such as ibis, spoonbills, egrets, herons, cormorants and pelicans can undertake rapid, long-distance flights (up to thousands of kilometres), including in some cases crossing to Papua New Guinea (McGinness et al. 2024). Many of the most important aggregation sites for these species, which can include dense congregations of hundreds to tens of thousands of birds, occur in the nationally significant and intensively managed Murray-Darling Basin (MDB), Australia’s largest river system located in south-east Australia (Kingsford et al. 2010). Therefore, given the risks to these sites and species, we established a new avian influenza surveillance and monitoring program in the MDB in 2025-26.

Animal ethics approval for bird monitoring and sampling was obtained from the CSIRO Wildlife, Large, and Laboratory Animal Ethics Committee (2025-33).

We conducted five fieldtrips in seven regions within the MDB to wetlands important for waterbirds between September 2025 and February 2026 (Figure 1). Figure 1 We collected 628 environmental faecal samples into 3mL of phosphate buffered gelatine saline viral transport (Table 1; Supplemental Methods). Of these, three were positive for influenza A virus (Table 1) but negative for H5 and H7 using qPCR (Supplemental Methods).

**Table 1.** Waterbird species targeted for environmental faecal sampling at important wetlands in the Murray-Darling Basin in spring/summer of 2025-26.

| Location | Dates | Species | Virus detection (qPCR) |
| --- | --- | --- | --- |
| Murrumbidgee – Western Lakes region (near Balranald, NSW) | Sep 13-14, 2025 | Australian Pelican *Pelecanus conspicillatus*<br>Black Swan *Cygnus atratus*<br>Grey Teal *Anas gracilis*<br>Mix 1*<br>Mix 2**<br>Little Black Cormorant *Phalacrocorax sulcirostris* | 0/5<br>0/10<br>**1/30**<br>0/17<br>**1/29**<br>0/8 |
| Murrumbidgee – Gayini (near Maude, NSW) | Sep 14-15, 2025 | Mix 3**<br>Mix 4*<br>Pied Stilt *Himantopus leucocephalus* | 0/22<br>0/19<br>0/14 |
| Macquarie Marshes (near Warren, NSW) | Oct 10-14, 2025 | Australian Pelican *Pelecanus conspicillatus*<br>Australian Shelduck *Tadorna tadornoides*<br>Australian White Ibis *Threskiornis moluccus*<br>Black Swan *Cygnus atratus*<br>Grey Teal *Anas gracilis*<br>Little Black Cormorant *Phalacrocorax sulcirostris*<br>Magpie Goose *Anseranas semipalmata*<br>Plumed Egret *Ardea plumifera*<br>Mix 2**<br>Unknown waterbird | 0/3<br>0/29<br>0/3<br>0/11<br>0/3<br>0/1<br>0/36<br>0/11<br>0/37<br>0/2 |
| Gwydir Wetlands (near Moree, NSW) | Nov 8-11, 2025 | Mix 2**<br>Mix 5**<br>Mix 6***<br>Yellow-billed Spoonbill *Platalea flavipes*<br>Straw-necked Ibis *Threskiornis spinicollis* | **1/7**<br>0/38<br>0/40<br>0/1<br>0/26 |
| Narran Lakes (near Brewarrina, NSW) | Nov 7-8, 2025 | Mix 2**<br>Mix 7°<br>Mix 8°°<br>Mix 9°°°<br>Mix 10\$<br>Pied Stilt *Himantopus leucocephalus* | 0/17<br>0/20<br>0/3<br>0/14<br>0/10<br>0/22 |
| Millewa Forest and Deniliquin Region (NSW) | Dec 9-13, 2025 | Mix 2**<br>Australian White Ibis *Threskiornis moluccus*<br>Plumed Whistling Duck *Dendrocygna eytoni*<br>Grey Teal *Anas gracilis* | 0/3<br>0/14<br>0/37<br>0/27 |
| Mid-Murray – Kerang (VIC) | Feb 3-4, 2026 | Mix 11\$§ | 0/59 |
| **GRAND TOTAL** |  |  | **3/628** |

\*Mix 1 - Grey Teal *Anas gracilis*, Pacific Black Duck *Anas superciliosa*
\*\*Mix 2 – Grey Teal *Anas gracilis*, Chestnut Teal *Anas castanea*, Pacific Black Duck *Anas superciliosa*
\*\*\*Mix 3 - Australian White Ibis *Threskiornis moluccus*, Royal Spoonbill *Platalea regia*, Yellow-billed Spoonbill *Platalea flavipes*
\*Mix 4 - Australian White Ibis *Threskiornis moluccus*, Yellow-billed Spoonbill *Platalea flavipes*
\*\*Mix 5 - Wandering Whistling Duck *Dendrocygna arcuata*, Freckled Duck *Stictonetta naevosa*, Pacific Black Duck *Anas superciliosa*, Grey Teal *Anas gracilis*, Chestnut Teal *Anas castanea*, Maned Duck *Chenonetta jubata*
\*\*\*Mix 6 – Little Egret *Egretta garzetta*, Plumed Egret *Ardea plumifera*, Great Egret *Ardea alba*, Australian White Ibis *Threskiornis moluccus*, Straw-necked Ibis *Threskiornis spinicollis*, Glossy Ibis *Plegadis falcinellus*, Royal Spoonbill *Platalea regia*, Yellow-billed Spoonbill *Platalea flavipes*
°Mix 7 - Grey Teal *Anas gracilis*, Chestnut Teal *Anas castanea*, Pacific Black Duck *Anas superciliosa*, Black Swan *Cygnus atratus*
°°Mix 8 – Australian White Ibis *Threskiornis moluccus*, Straw-necked Ibis *Threskiornis spinicollis*, Glossy Ibis *Plegadis falcinellus*, Royal Spoonbill *Platalea regia*, Yellow-billed Spoonbill *Platalea flavipes*
°°°Mix 9 - Glossy Ibis *Plegadis falcinellus*, Plumed Egret *Ardea plumifera*, Great Egret *Ardea alba*
\$Mix 10 - Australian White Ibis *Threskiornis moluccus*, Pacific Heron *Ardea pacifica*, Little Egret *Egretta garzetta*, Great Egret *Ardea alba*, Royal Spoonbill *Platalea regia*
\$§Mix 11 – Grey Teal *Anas gracilis* & Black Swan *Cygnus atratus*

**Figure 1.**
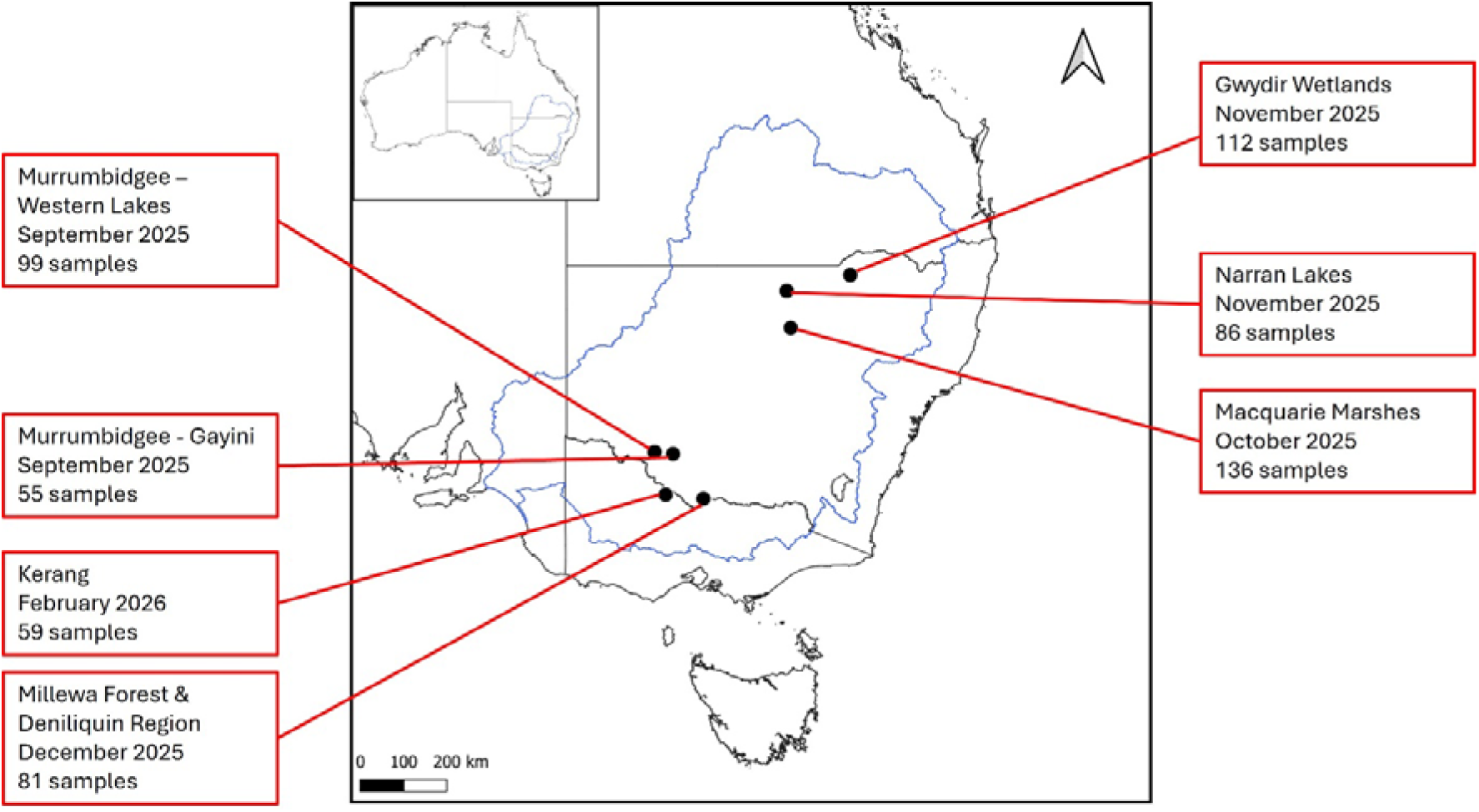
Overview of waterbird surveillance at wetlands of the Murray-Darling with major waterbird aggregations between September 2025 and February 2026. Blue outline shows the Murray-Darling Basin. Black lines show Australian state borders.

We attempted full genome sequencing on the three positive samples. Of these, we were able to sequence a complete H10N8 genome from a mixed sample group containing Grey Teal (*Anas gracilis*), Chestnut Teal (*Anas castanea)* and Pacific Black Duck (*Anas superciliosa)* sampled in the Murrumbidgee Western Lakes region near Balranald, NSW (∼34.617°S, 143.567°E) in September 2025 and a partial H10N7 genome from mixed sample group of Grey Teal, Chestnut Teal and Pacific Black Duck sampled in the Gwydir Wetlands region near Moree, NSW (∼29.464° S, 149.845° E) in November 2025. Genomes have been deposited in GenBank (Accession PZ736887-PZ736899).

These genomes are reassorted, but share the HA, NP and NS segments (Figure 2, S1, S2). While all sequences in the H10N7 genome were mostly closely related to sequences from Australian wild birds, some segments of the H10N8 genome fell into clades that are poorly sampled in Australia. Specifically, the PB2, PA, NA from the H10N8 genome were sister to only a single Australian sequence, nestled within a clade of Eurasian genomes (Figure 2, S1, S2), as opposed to being nestled within a clade with >10 Australian genomes. The M sequence from this H10N8 genome was not sister to any Australian sequences, and the maximum likelihood tree had a long branch between the sequence generated in this study and sister sequences, suggesting undersampled diversity. This is not surprising as, since 2022, only select sequences from Australian ducks (H10, H5, H7) and shorebirds (H3, H4) have been deposited in GenBank (e.g., see Wille et al. 2024a; Wille et al. 2024b).

**Figure 2.**
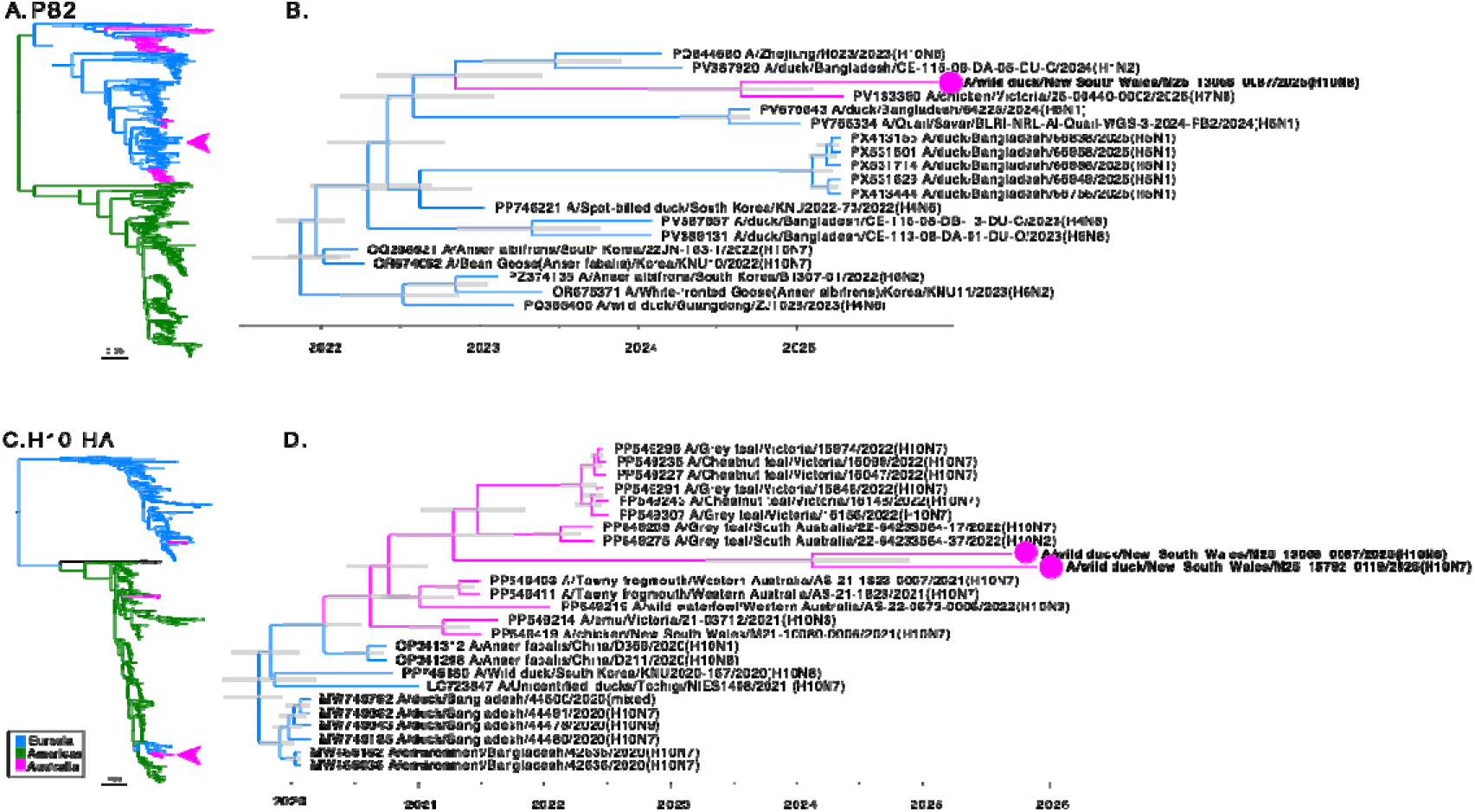
Phylogenetic trees for the PB2 and H10 sequences generated in this study. (A, C) Maximum likelihood trees of a representative set of global diversity. Trees were mid-point rooted, illustrating the deep phylogenetic division between viruses circulating in Eurasia and the Americas. Scale bar is number of substitutions per site and arrows denoted position of sequences generated in this study. (B, D) Excerpt of time structed phylogenetic trees, scale bar indicates time in years, and viruses are denoted by filled circles. As the PB2 sequence of the H10N7 genome is partial (∼500bp), is has not been included here. The top blast hit is shown in Table S1. Tree files are available at https://github.com/michellewille2/MDB_surveillance/ and trees for the internal segments and NA sequences are in Figure S1-S2.

During field trips we also conducted monitoring to document local species composition and look for signs of sick or dead birds. This included 19 stationary point surveys each lasting 20 minutes across the six locations visited, during which we recorded 4,539 of 54 waterbird species. We also conducted 25 drone searches lasting 5-15 minutes each across four locations to view inaccessible areas.

During monitoring we encountered one apparently unhealthy Australasian Shoveller (*Anas rhynchotis)* and took oropharyngeal and cloacal samples, which were negative for influenza A virus by qPCR. We also encountered three dead waterbirds – two Australian Pelican (*Pelecanus conspicillatus)* and one Australasian White Ibis (*Threskiornis moluccus)*, all too decomposed for sampling.

There is growing evidence that collecting as much monitoring and surveillance data as possible from wild birds is critical to disease impact mitigation and species conservation (e.g., see Knief et al. 2026). Establishment of this surveillance and monitoring program at large inland wetlands of the MDB involved significant logistical effort, as natural wetlands with waterbird aggregations in the MDB differ significantly from urban wetlands and coastal migratory bird aggregation sites where most prior surveillance activities for wild Australian birds were conducted. In MDB wetlands, target species may spend most or all of daylight hours either standing in shallow water or swimming, making faecal samples inaccessible. At times this posed a significant challenge during our fieldwork. Target species may also aggregate in areas far from the shoreline requiring a boat or kayak for access, also a situation that occurred regularly during our fieldwork. Implementation of our program, including detection of multiple low pathogenicity strains of influenza A, demonstrates that monitoring and surveillance can successfully be completed in this large inland wetland setting. This, combined with publication of full genome sequencing, helps to address undersampling of locations and species. Maintaining and enhancing this capability remains critical for early detection and rapid response in the case of an extensive inland outbreak.

## Funding Acknowledgement

This work was supported by the Australian Government through the Commonwealth Environmental Water Holder, part of the Department of Climate Change, Energy, the Environment and Water, with H5 bird flu funding. Michelle Wille is supported by an Australian Research Council Future Fellowship, and the WHO Collaborating Centre for Reference and Research on Influenza is supported by the Australian Centre for Disease Control.

## Author Acknowledgements

The authors would like to thank land holders, Traditional Owners, NSW DCCEEW, wetland and water managers in the Murray–Darling Basin, including NSW NPWS and NCCMA, and academic colleagues for facilitating access to wetlands and supporting planning and logistics for monitoring, surveillance and bird tracking activities. We are very grateful for their assistance and support. Further, we would like to acknowledge the assistance of Wildlife Health Australia for providing technical guidance and support from the National Avian Influenza Wild Bird surveillance program and connecting the project team with jurisdictional surveillance partners, and state governments and veterinary officers, CSIRO Australian Centre for Disease Preparedness (ACDP), state testing labs and academic colleagues for their input into monitoring and surveillance methodology and facilitation of sample testing.

