## Supplemental Methods for "Establishment of an avian influenza surveillance program in Australia’s largest river basin"

**Supplemental Materials**


| | |
|---|---|
| Supplemental Methods | 2 |
| Figure S1. Phylogenetic trees for segments PB1, PA, NP, M, NS | 4 |
| Figure S2. Phylogenetic trees for N7 and N8 | 5 |
| Table S1. Top blast hits for all segments | 6 |

**Supplemental Methods**

*Sample collection and screening*

Prior to sampling birds were surveyed from a distance using binoculars and/or a spotting telescope and identified to species level. For faecal sampling, we aimed to take a minimum of 30 faecal deposits per site. Where possible we took 30+ samples per species per site and at least 60 swabs total if we found sufficiently high numbers of birds (but avoiding oversampling if there were low numbers). If we encountered lower numbers of a species known to be particularly susceptible to carrying or suffering from the virus (particularly ducks or shorebirds), we took samples if 5-10 per species could be obtained. Only very fresh faecal samples (visible moisture) were collected from the ground, nests, or roosts. Whenever possible sampling was conducted in the early morning to limit UV damage. Samples were collected using 7.5cm wood-stemmed sterile cotton-tipped applicators (Multigate Medical Products), placed into 3mL of phosphate buffered gelatine saline viral transport medium (VTM). Swabs were kept at 4°C for up to 7 days, following storage at -80°C.

Samples collected in New South Wales (NSW) were tested at the Elizabeth Macarthur Agricultural Institute and samples collected in Victoria were tested at the Victorian Department of Energy, Environment and Climate Action (DEECA), Agriculture Science and Technology.

For samples collected in NSW, RNA was extracted from 50µl of sample in VTM using the MagMax 96 Viral RNA Isolation Kit (Applied Biosystems, Thermo Fisher Scientific, AMB18365) using the Kingfisher 96 magnetic particle processor (Thermo Fisher Scientific, CAT 5400500). The RNA was tested using the AIV Type A RT-qPCR assay which targets the highly conserved matrix (M) gene (Heine et al. 2007). RT-qPCR reactions were performed using the AgPath-ID One-Step RT-PCR kit (Thermo Fisher Scientific, AM1005) on the QuantStudio 5 Real-Time PCR system (Thermo Fisher Scientific, A28133).

For samples collected in Victoria, RNA was extracted using the MagMax 96 Viral Isolation Kit (Applied Biosystems, Thermo Fisher Scientific) using the Kingfisher Flex (Thermo Fisher Scientific). The RNA was tested using the AIV Type A real-time hydrolysis probe (TaqMan®) based RT-qPCR assay, which targets the highly conserved matrix (M) gene [REF]. RT-qPCR reactions were performed using the AgPath-ID™ One-Step RT-PCR kit (Thermo Fisher Scientific).

*Virus Characterisation*

As NSW was the only state in which samples were positive, full genome sequencing was undertaken at EMAI. Amplicons for influenza A viruses were generated using previously published MB-Tuni primers (Zhou and Wentworth 2012; Zhou et al. 2009) with the SuperScript III One-Step RT-PCR system (ThermoFisher Scientific). PCR products were visualised on a 1% agarose gel, before purification using Agencourt AMPure beads (Beckman Coulter Life Sciences, A63881). Quantitation was undertaken using the Quant-iT dsDNA Assay kit, high sensitivity kit (Thermo Scientific, Q33120) on the Qubit 4 Fluorometer (Thermo Fisher

Scientific, Q33238). dsDNA end repair was completed using NEB next ultra II end repair kit (New England Biolab, E7546L).

Libraries were prepared using 200fmol of repaired cDNA and barcoded using Native Barcoding kit 96 v14 (Oxford Nanopore Technologies, SQK-NBD114.96). Libraries were run on MinION Flow Cell-DNA (Oxford Nanopore Technologies, FLO-MIN114) using a MinION Mk1B (Oxford Nanopore Technologies) for 8 h.

Genomes were assembled through a manually generated influenza virus library using the IRMA: Iterative Refinement Meta-Assembler Pipeline (Shepard et al. 2016). Quality control, such as the removal of short, low quality, and chimeric reads was automated through the IRMA pipeline. Coverage and depth statistics were generated using SAMtools (Danecek et al. 2021).

For all segments, two maximum likelihood trees were constructed. First, "global" trees using backbones reported in Wille et al. (2024) to demonstrate phylogenetic placement relative to globally circulating viruses. Second, "local" trees comprising only the top 20 blast hits, retrieved in Geneious Prime (6 July 2025), as well as a North American outgroup sequence retrieved from the BV-BRC database (https://www.bv-brc.org/). In both cases, maximum likelihood trees incorporating the best-fit model of nucleotide substitution were estimated using IQ-Tree (Nguyen et al. 2015), and 1000 ultrafast bootstraps. Trees were visualised using Fig Tree v1.4. For the HA and PB2, we performed a linear regression of root-to-tip distances against year of sampling with TempEst (Rambaut et al. 2016). Subsequently, a time-scaled phylogenetic tree was estimated using BEAST v1.10.4 (Drummond et al. 2012), under the uncorrelated lognormal relaxed clock (Li and Drummond 2012) and SRD06 codon structured nucleotide substitution model (Shapiro et al. 2006), and the Bayesian skyline coalescent tree prior (Drummond et al. 2005). One hundred million generations were performed, and convergence was assessed using Tracer v1.8 (http://tree.bio.ed.ac.uk/software/tracer/). A maximum credibility lineage tree was generated using TreeAnnotator following the removal of 10% burn-in, and were visualised using Fig Tree v1.4 (http://tree.bio.ed.ac.uk/software/figtree/).

*Waterbird monitoring*

For point surveys we recorded all waterbirds we could see from a stationary point using a 20-60 x 82 spotting scope to species level. To estimate the number of birds per unit area (to indicate density and for potential comparison between sites), we noted how many birds were inside either a 360° circle with a radius of 80m or a 180° half circle with a radius of 160m, depending on wetland configuration: if we were surveying from the middle of a wetland we used the full circle technique; if we were surveying from a shoreline into a wetland we used the half circle technique. We recorded all waterbirds to species level and in the following categories: i) [apparently] healthy, sick or dead; ii) adult or juvenile; iii) at nest or not at nest. We limited counts to 20 minutes and recorded the time, surveyor, scribe, and weather conditions.

Where permissions could be obtained and when weather conditions were suitable, we also conducted drone searches to complement point-count surveys. The primary purpose of the drone searches was to search for sick or dead birds that were not visible from accessible ground survey points. Drone flights generally lasted 5-15 minutes and a video of the entire flight was recorded. We noted the approximate area covered on a printed map.

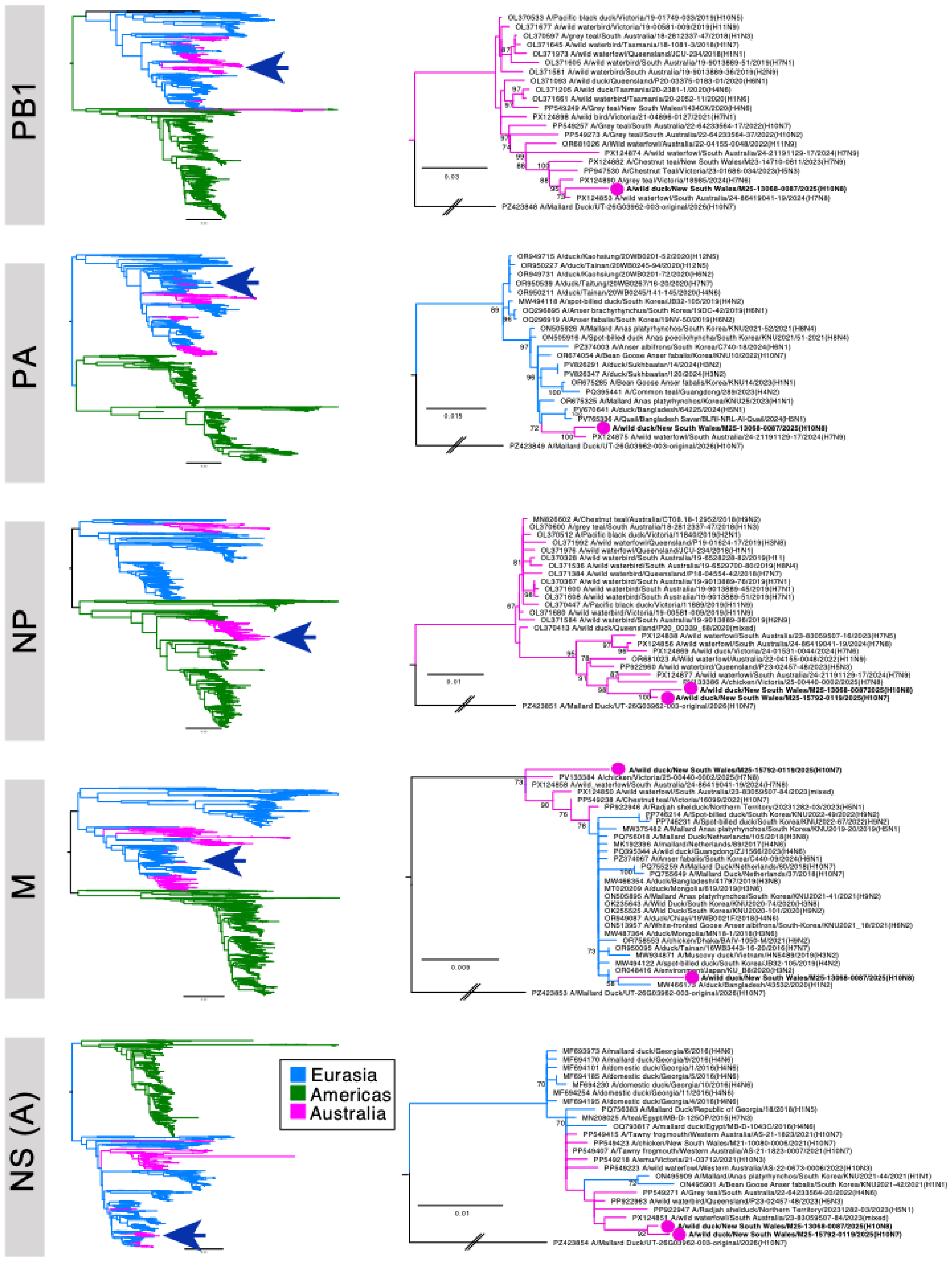


*Figure S 1. Phylogenetic trees for segments PB1, PA, NP, M, NS, for all viruses sequenced in this study. Viruses are denoted by arrows (on the left panels) and a filled circle (on the right panels). Trees were rooted based on the geographic division between American and Eurasian clades, with trees on the right rooted using A/Mallard Duck/UT-26G03962-003-original/2026(H10N7). Branches are coloured based on geographic region. The scale bar indicates the number of substitutions per site.*

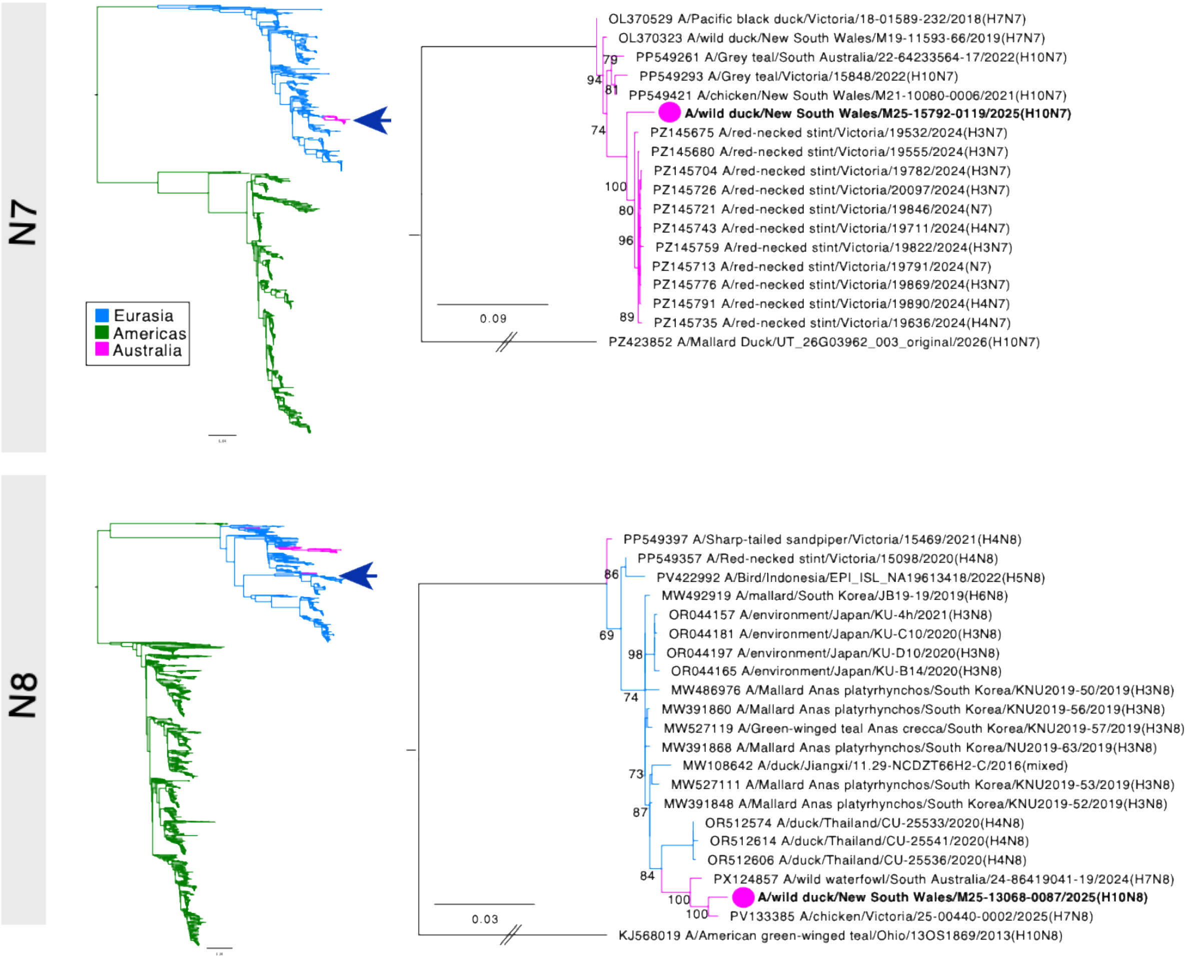


*Figure S 2. Phylogenetic trees for the N7 and N8 subtype of the NA segment. Viruses are denoted by arrows (on the left panels) and a filled circle (on the right panels). Trees were rooted based on the geographic division between American and Eurasian clades, with trees on the right rooted using a North American clade N7 or N8 sequence. Branches are coloured based on geographic region. The scale bar indicates the number of substitutions per site.*

| | M25-13086-0087 | | | M25-15792-0119 | | |
|---|---|---|---|---|---|---|
| PB 2 | A/chicken/Victoria/25-00440-0002/2025(H7N8) | PV133390 | 98.90% | A/wild waterfowl/South Australia/23-83059507-80/2023(H7N5) | PX124905 | 98.70% |
| | A/Spot-billed duck/South Korea/KNU2022-73/2022(H4N6) | PP746221 | 97.90% | A/Chestnut teal/New South Wales/M23-14710-0811/2023(H7N9) | PX124881 | 98.70% |
| | A/Anser albifrons/South Korea/22JN-163-1/2022(H10N7) | OQ296821 | 97.90% | A/wild waterfowl/South Australia/23-83059507-16/2023(H7N5) | PX124834 | 98.70% |
| | A/Bean Goose(Anser fabalis)/Korea/KNU10/2022(H10N7) | OR674052 | 97.80% | A/chicken/New South Wales/M21-10080-0006/2021(H10N7) | PP549416 | 98.70% |
| | A/Anser albifrons/South Korea/22MC-41/2022(H6N2) | OQ296845 | 97.70% | A/Chestnut teal/Victoria/16148/2022(H10N7) | PP549240 | 98.70% |
| PB 1 | A/wild waterfowl/South Australia/24-86419041-19/2024(H7N8) | PX124853 | 98.90% | | | |
| | A/grey teal/Victoria/18985/2024(H7N6) | PX124890 | 98.80% | | | |
| | A/Chestnut Teal/Victoria/23-01686-034/2023(H5N3)) | PP947530 | 98.20% | | | |
| | A/Chestnut teal/New South Wales/M23-14710-0811/2023(H7N9) | PX124882 | 98.00% | | | |
| | A/Wild waterfowl/Australia/22-04155-0048/2022(H11N9) | OR681026 | 97.60% | | | |
| PA | A/wild waterfowl/South Australia/24-21191129-17/2024(H7N9) | PX124875 | 99.20% | | | |
| | A/Mallard(Anas platyrhynchos)/Korea/KNU25/2023(H1N1) | OR675325 | 98.50% | | | |
| | A/Mallard(Anas platyrhynchos)/South Korea/KNU2021-52/2021(H8N4) | ON505926 | 98.40% | | | |
| | A/Bean Goose(Anser fabalis)/Korea/KNU10/2022(H10N7) | OR674054 | 98.40% | | | |
| | /Spot-billed duck(Anas poecilohyncha)/South Korea/KNU2021-51/2021(H8N4) | ON505916 | 98.20% | | | |
| HA | A/Grey teal/South Australia/22-64233564-37/2022(H10N2) | PP549275 | 96.50% | A/Tawny frogmouth/Western Australia/AS-21-1823/2021(H10N7) | PP549411 | 96.40% |
| | A/Tawny frogmouth/Western Australia/AS-21-1823/2021(H10N7) | PP549411 | 96.40% | A/Grey teal/South Australia/22-64233564-37/2022(H10N2) | PP549275 | 96.30% |
| | A/Grey teal/South Australia/22-64233564-17/2022(H10N7) | PP549259 | 96.40% | A/Tawny frogmouth/Western Australia/AS-21-1823-0007/2021(H10N7) | PP549403 | 96.30% |
| | A/Tawny frogmouth/Western Australia/AS-21-1823-0007/2021(H10N7) | PP549403 | 96.30% | A/Grey teal/South Australia/22-64233564-17/2022(H10N7) | PP549259 | 96.30% |
| | A/Grey teal/Victoria/15848/2022(H10N7) | PP549291 | 96.30% | A/Eurasian teal/South Korea/JB32-15/2019(H10N7) | MW494148 | 96.20% |
| NP | A/wild waterbird/Queensland/P23-02457-48/2023(H5N3) | PP922960 | 98.10% | A/wild waterbird/Queensland/P23-02457-48/2023(H5N3) | PP922960 | 98.50% |
| | A/wild waterfowl/South Australia/24-21191129-17/2024(H7N9) | PX124877 | 98.00% | A/wild waterfowl/South Australia/24-21191129-17/2024(H7N9) | PX124877 | 98.40% |
| | A/chicken/Victoria/25-00440-0002/2025(H7N8) | PV133386 | 97.80% | A/Wild waterfowl/Australia/22-04155-0048/2022(H11N9) | OR681023 | 98.10% |
| | A/Wild waterfowl/Australia/22-04155-0048/2022(H11N9) | OR681023 | 97.70% | A/chicken/Victoria/25-00440-0002/2025(H7N8) | PV133386 | 98.10% |

| | | | | | | |
|---|---|---|---|---|---|---|
| | A/wild waterfowl/South Australia/23-83059507-16/2023(H7N5) | PX124838 | 97.30% | A/wild waterfowl/South Australia/24-86419041-19/2024(H7N8) | PX124856 | 97.70% |
| NA | A/chicken/Victoria/25-00440-0002/2025(H7N8) | PV133385 | 99.10% | A/red necked stint/Victoria/19636/2024(H4N7) | PZ145735 | 97.00% |
| | /wild waterfowl/South Australia/24-86419041-19/2024(H7N8) | PX124857 | 98.50% | A/red necked stint/Victoria/19532/2024(H3N7) | PZ145675 | 97% |
| | A/Mallard(Anas platyrhynchos)/South Korea/KNU2019-52/2019(H3N8) | MW391848 | 97.70% | A/red necked stint/Victoria/19555/2024(H3N7) | PZ145680 | 97% |
| | A/Mallard(Anas platyrhynchos)/South Korea/KNU2019-53/2019(H3N8) | MW527111 | 97.60% | A/red necked stint/Victoria/19791/2024(N7) | PZ145713 | 96.90% |
| | A/Mallard(Anas platyrhynchos)/South Korea/NU2019-63/2019(H3N8) | MW391868 | 97.50% | A/red necked stint/Victoria/19846/2024(N7) | PZ145721 | 96.90% |
| M | A/environment/Japan/KU-B8/2020(H3N2) | OR048416 | 99.20% | A/wild waterfowl/South Australia/24-86419041-19/2024(H7N8) | PX124858 | 99.10% |
| | A/Radjah shelduck/Northern Territory/20231282-03/2023(H5N1) | PP922946 | 99.10% | A/chicken/Victoria/25-00440-0002/2025(H7N8) | PV133384 | 98.90% |
| | A/duck/Chiayi/19WB0021F/2018(H4N6) | OR949087 | 99.10% | A/Mallard Duck/Netherlands/37/2018(H10N7) | PQ755649 | 98.70% |
| | A/White-fronted Goose(Anser albifrons)/South Korea/KNU2021-18/2021(H6N2) | ON513957 | 99.10% | A/Radjah shelduck/Northern Territory/20231282-03/2023(H5N1) | PP922946 | 98.70% |
| | A/Mallard(Anas platyrhynchos)/South Korea/KNU2021-41/2021(H9N2) | ON505895 | 99.10% | A/duck/Chiayi/19WB0021F/2018(H4N6) | OR949087 | 98.70% |
| NS | A/wild waterfowl/South Australia/23-83059507-84/2023(mixed) | PX124851 | 99.20% | A/wild waterfowl/South Australia/23-83059507-84/2023(mixed) | PX124851 | 99.00% |
| | A/wild waterbird/Queensland/P23-02457-48/2023(H5N3) | PP922963 | 99.20% | A/wild waterbird/Queensland/P23-02457-48/2023(H5N3) | PP922963 | 99.00% |
| | A/Tawny frogmouth/Western Australia/AS-21-1823-0007/2021(H10N7) | PP549407 | 99.20% | A/Tawny frogmouth/Western Australia/AS-21-1823-0007/2021(H10N7) | PP549407 | 99.00% |
| | A/Tawny frogmouth/Western Australia/AS-21-1823/2021(H10N7) | PP549415 | 99.00% | A/Tawny frogmouth/Western Australia/AS-21-1823/2021(H10N7) | PP549415 | 98.90% |
| | (A/Mallard(Anas platyrhynchos)/South Korea/KNU2021-44/2021(H1N1) | ON495909 | 99.00% | A/Mallard(Anas platyrhynchos)/South Korea/KNU2021-44/2021(H1N1) | ON495909 | 98.90% |

*Table S 1. Top blast hits for all segments*
